# Spatial transcriptomic atlas of shoot organogenesis in tomato callus

**DOI:** 10.1101/2023.02.24.529793

**Authors:** Xiehai Song, Pengru Guo, Meiling Wang, Lichuan Chen, Jinhui Zhang, Mengyuan Xu, Naixu Liu, Min Liu, Liang Fang, Xun Xu, Ying Gu, Keke Xia, Bosheng Li

**Affiliations:** Peking University Institute of Advanced Agricultural Sciences, Shandong Laboratory of Advanced Agriculture Sciences in Weifang, Weifang, Shandong 261325, China; BGI-Shenzhen, Shenzhen, Guangdong 518083, China; Department of Biology, School of Life Sciences, Southern University of Science and Technology, Shenzhen, Guangdong 518005, China; BMKMANU, Qingdao, Shandong 266500, China

**Keywords:** shoot regeneration, cellular reprogramming, spatial transcriptome, callus, vascular tissue

## Abstract

Callus is a reprogrammed transitional cell mass during plant regeneration. Pluripotent callus cells develop into fertile shoots through *de novo* shoot organogenesis. This study represents a pioneering effort in exploring the spatial transcriptome of tomato callus during shoot regeneration, using technologies including BGI Stereo-seq, BMKMANU S1000, and 10x Visium. The results indicate that the callus comprises highly heterogeneous cells, classified into various cell types based on spatial gene expression and histological observation, including epidermis, shoot primordium, vascular tissue, inner callus, and outgrowth shoots. The developmental trajectories from shoot primordium to outgrowth shoot are traced, and vascular tissue development is characterized. The single-cell resolution spatial approach reveals the origin of shoot primordia from the sub-epidermis. The spatial full length RNA sequencing shows high incompletely spliced (IS) ratios in the shoot primordium cells. These findings enhance our knowledge of plant organogenesis and highlight the significance of spatial biology in plant research.

## Introduction

Plant regeneration, the ability of injured plants to repair or reconstruct new tissues or organs, is determined by the totipotency or pluripotency of plant cells (Ikeuchi et al., 2019). There are various modes of regeneration in higher plants, including tissue repair, *de novo* organogenesis, and somatic embryogenesis (Birnbaum and Sánchez Alvarado, 2008; Ikeuchi et al., 2019). In particular, *de novo* organogenesis involves the formation of new shoots or roots from established shoot or root apical meristems, known as *de novo* shoot organogenesis (DNSO) or *de novo* root organogenesis (DNRO), respectively (Radhakrishnan et al., 2018; Shin et al., 2020; Xu, 2018). The in vitro tissue culture technique, which involves incubating various explants in a nutrient-rich medium supplemented with different concentrations of auxin and cytokinin, has been developed and applied in many plant species (Ikeuchi et al., 2016; Ikeuchi et al., 2013; Skoog and Miller, 1957). This method generates stable transgenic plants with desirable traits through *Agrobacterium* in crop breeding. However, it has limitations in essential crops like soybean, wheat, and maize (Que et al., 2014; Wang et al., 2022; Xu et al., 2022). Tomato (*Solanum lycopersicum*.), an economically significant crop cultivated globally, has the tissue culture highly dependent on its genotype (Bhatia et al., 2004; Praveen and Nanna, 2011). Thus, understanding the *de novo* organogenesis process and the underlying molecular mechanism is crucial and of great value for improving tissue culture technology in crops.

DNSO has been extensively studied in the genetically tractable model plant *Arabidopsis thaliana* in recent years. When root or hypocotyl explants are placed in the auxin-rich callus induction medium (CIM), the xylem-pole pericycle cells are reprogrammed to form the pluripotent callus (Atta et al., 2009). This pluripotent callus is crucial for DNSO and resembles the root primordium, with key regulators of root meristem formation, such as *WUSCHEL-RELATED HOMEOBOX5* (*WOX5*), *WOX7, PLETHORA 1* (*PLT1*), *PLT2, LATERAL ORGAN BOUNDARIES-DOMIAN 16* (*LBD16*), *SHORT ROOT* (*SHR*), and *SCARECROW* (*SCR*) playing vital roles in inducing pluripotency (Okushima et al., 2007; Shin et al., 2020; Sugimoto et al., 2010). A subpopulation of pluripotent callus cells then acquires shoot identity and develops into shoot apical meristem (SAM) and entire shoots upon incubation in the cytokinin-rich shoot-inducing medium (SIM) (Shin et al., 2020). DNSO from callus involves a series of cell fate determination, transformation, and development. Early promoters of DNSO include the *CUP-SHAPED COTYLEDON 1* (*CUC1*) and *CUC2* transcription factors and the auxin efflux transporter PIN1, whose spatial locations in SIM determine the future sites of shoot primordia/progenitors (Gordon et al., 2007; Varapparambath et al., 2022). Upstream regulators of DNSO, such as the *APETALA2/Ethylene Responsive Factor* (*AP2/ERF*)-type transcription factors *ENHANCER OF SHOOT REGENERATION 1* (*ESR1*) and its paralog *ESR2, PLT3, PLT5*, and *PLT7*, activate the expression of *CUC* genes (Daimon et al., 2003; Kareem et al., 2015; Matsuo et al., 2011). A key molecular event in DNSO is the selective transcriptional activation of *WUSCHEL* (*WUS*), triggered by the cytokinin-responsive B-type ARRs (*ARR1, ARR10*, and *ARR12*) and the class III homeodomain-leucine zipper (HD Zip III) transcription factors *PHABULOSA* (*PHB*), *PHAVOLUTA* (*PHV*), and *REVOLUTA* (*REV*) (Zhang et al., 2017; Zubo et al., 2017). *WUS-*expressing cells mark the shoot primordia/progenitors, and organized *WUS* expression is critical for further SAM formation (Gordon et al., 2007). Other shoot meristem regulators, such as *SHOOT MERISTEMLESS* (*STM*) and *CLAVATA3* (*CLV3*), are also expressed and functional in this process (Gordon et al., 2007; Shin et al., 2020). While the molecular mechanism of DNSO has been well studied in *Arabidopsis*, it remains elusive in tomato, especially in terms of single-cell and spatial analysis.

DNSO from callus involves the selection of shoot primordia/progenitors from a mass of pluripotent callus, followed by a series of cell fate transformations and development, and the spatial expression of numerous shoot promoters. This process highlights the heterogeneity of the callus and the importance of its spatial information. The shoot progenitor cell, for example, is determined through the expression of *WUS* in sporadic cells in the SIM-cultured callus, emphasizing the significance of the location of *WUS* expression. (Gordon et al., 2007; Varapparambath et al., 2022; Zhang et al., 2017). However, the determination of the spatial expression pattern of *WUS* and the transcriptomic differences between shoot progenitors and the surrounding callus cells are currently unknown.

Advances in single-cell/single-nuclear RNA sequencing (sc/snRNA-seq) techniques such as 10x genomics Chromium Single Cell and BGI DNBelab C4 allow for expression pattern profiling of heterogeneous tissues and organs at a single-cell level in both animals and plants. Furthermore, spatial transcriptome techniques such as BGI Stereo-seq and 10x Visium have been used to resolve the in situ spatial transcriptomes of tissues, where the location information of transcripts is critical (Liu et al., 2022; Xia et al., 2022). These emerging tools can potentially investigate comprehensive molecular and cellular events in DSNO. For instance, a recent study applied sc-RNA-seq to *Arabidopsis* callus, revealing the ability of its middle cell layers to regenerate shoots and identifying a related mechanism involving the promotion of auxin production and the enhancement of cytokinin sensitivity (Zhai and Xu, 2021). This study primarily focuses on the specific WOX5/7 factors and their regulation mechanism, with limited genome-wide discoveries on a large scale. Additionally, the *Arabidopsis* hypocotyl callus used in this research is different from the callus used in agricultural breeding for transgene insertion.

In this study, three spatially resolved transcriptomic profiling techniques were used to analyze tomato callus during shoot regeneration: BGI Stereo-seq, BMKMANU S1000, and 10x Visium. The main cell types of the callus were identified through histological observation and specific marker genes. The developmental trajectories of shoot primordia to outgrowth shoots were reconstructed. Additionally, the characterization of marker genes in vascular tissue revealed the diversity of its development and function. The origin of shoot primordia was investigated using the single-cell resolution BGI Stereo-seq, while the spatial alternative splicing was studied based on the BMKMANU S1000 approach.

## Results

### Spatially and single-nuclear resolved transcriptomic atlas of shoot regeneration from tomato callus

The process of DNSO in plants typically includes pluripotency acquisition in callus, selection of shoot progenitors/shoot primordia, the establishment of the SAM, and shoot outgrowth. To gain a thorough molecular understanding of this complex process, we performed spatial transcriptomic profiling on tomato callus grown in a regeneration medium, when notable regenerated shoots emerged. We utilized three different spatial transcriptome sequencing techniques, the BGI single-cell Stereo-seq (ScStereo-seq), the BMKMANU S1000, and the 10x Visium, to obtain the spatial transcriptome data of the samples. Additionally, we performed single-nuclear transcriptome profiling of the tomato callus at the same time to capture a broader range of cell types (Figure 1A). The samples are listed in Figure 1B. ScStereo-seq generated single-cell data based on a cell mask that contained cell boundary information obtained from cell wall-staining images (Figure 1B - Sample i). After eliminating low-quality cells, data from 6,759 single cells with an average of 1,161 unique molecular identifier (UMI) and 608 genes per cell were obtained. Unsupervised clustering analysis of these cells was conducted by the R package Seurat, resulting in 8 clusters (Cluster i, Supplemental Table 1) that were spatially distributed on the cell mask (Figure 1C). BMKMANU S1000 captured 3,511 spots (50μm resolution) with an average of 1,388 UMI and 990 genes per spot in Sample ii, while the 10x Visium captured 2,051 spots (100μm resolution) with an average of 7,020 UMI and 2,797 genes per spot in Sample iii. The spots were analyzed using Seurats unsupervised clustering analysis, resulting in 10 clusters (Cluster ii, Supplemental Table 2) in Sample ii and 12 clusters (Cluster iii, Supplemental Table 3) in Sample iii, which were spatially distributed in the samples, respectively (Figure 1D and 1E).

**Figure 1.**
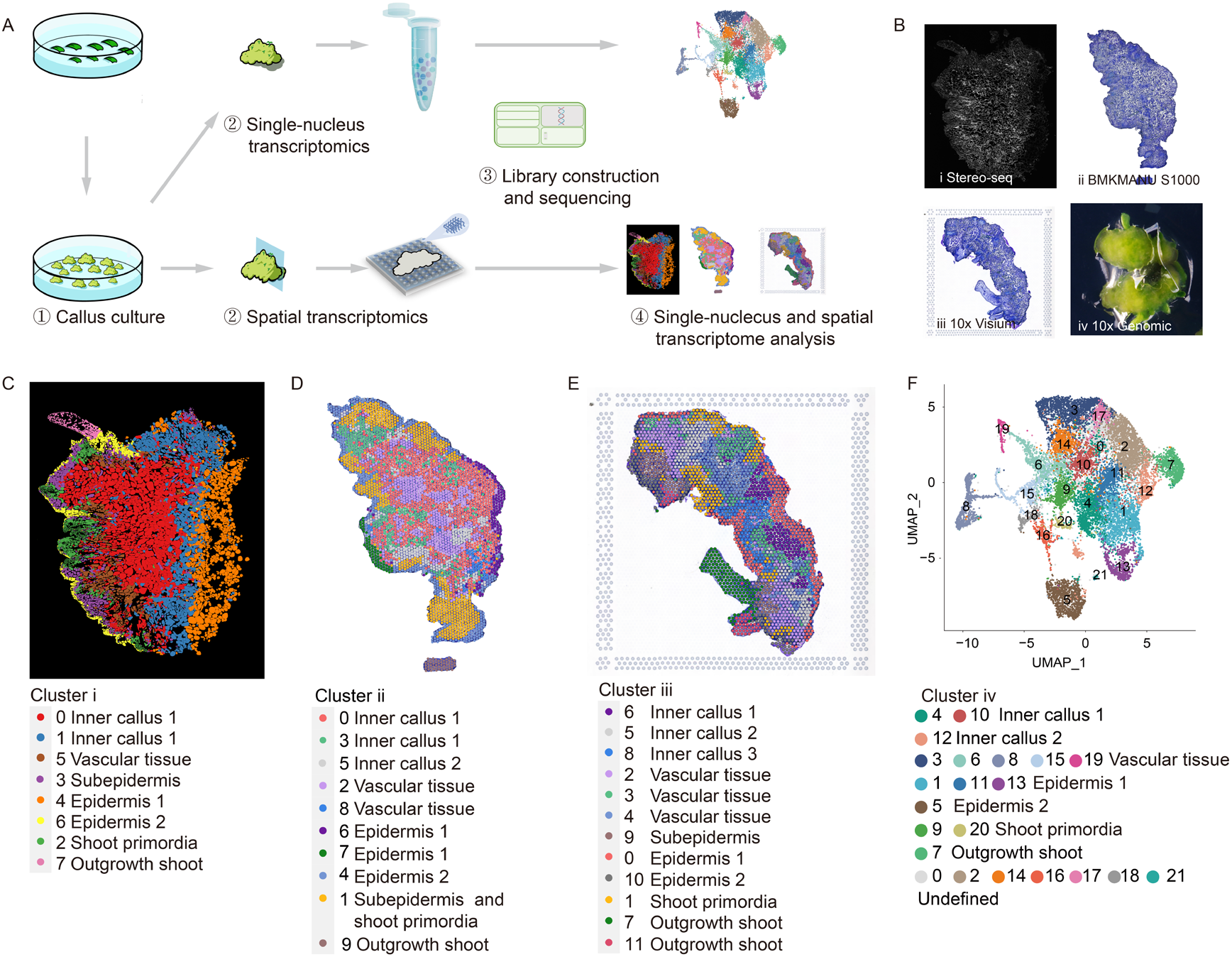
Cell clustering and cell-type identification in Tomato Callus based on Spatial and Single-nuclear Transcriptomics. **(A)** Spatial and single-nuclear RNA sequencing workflow in tomato callus samples cultured in a cytokinin-rich regeneration medium when regenerated shoots emerged. **(B)** The samples were subjected to different approaches, as indicated. Sample i, ii, iii, iv were used for Stereo-seq, BMKMANU S1000, 10x Visium, and 10x Chromium Single Cell, respectively. Sample i, ii, and iii are tissue sections with 10 μm thickness. Sample i was treated with Fluorescent Brightener 28 (FB) for cell wall staining, and Samples ii and iii were treated with Toluidine blue (TB) staining to obtain histological information. **(C-F)** Visualization of identified cell types in Sample i by the Stereo-seq (C), in Sample ii by BMKMANU S1000 (D), in Sample iii by 10x Visium (E), and in Sample iv by 10x Chromium Single Cell (F). The clusters identified by different approaches are named Cluster i, ii, iii, and iv, respectively, and the corresponding markers identified in different clusters are listed in Supplemental Table 1-4. Cell types identified by different platforms are marked by colors and defined consistently, and their correlations are shown in Supplemental Figure 1

The histological observation and the expression of specific marker genes were analyzed to identify the cell types represented by each cluster. Seven hundred fifty-one type-1 epidermal cells (Cluster i4, epidermis 1), 1,016 shoot primordia cells (Cluster i2), 599 vascular tissue cells (Cluster i5), and 214 shoot outgrowth cells (Cluster i7) were annotated based on typical marker genes for cell types in *Arabidopsis*, such as *PROTODERMAL FACTOR1* (*PDF1*), *A. thaliana MERISTEM LAYER 1* (*ATML1*), *3-ketoacyl-CoA synthase 11* (*KCS11*) in Epidermis 1 cells, *photosystem I light harvesting complex gene 2* (*LHCA2*), *chlorophyll A/B binding protein 1* (*CAB1*), *Rubisco small subunit 3B* (*RBCS3B*) in outgrowth shoot cells, *CUC1, WUS, STM* in shoot primordia cells, and *SMAX1-like 3* (*SMXL3*), *ATP-binding cassette G14* (*ABCG14*) in vascular tissue cells (Supplemental Figure 1). Meanwhile, 2,629 inner type-1 callus cells (Cluster i0 and i1, inner callus 1), 1,002 sub-epidermis cells (Cluster i3), and 548 epidermis 2 cells (Cluster i6) were annotated by their spatial locations (Figure 1C).

We analyzed the relationship between Clusters i and ii by comparing their cell types and found a strong correlation (Supplemental Figure 1). Based on this correlation, spots in Cluster ii were categorized into various cell types, including 1,445 inner callus 1 cells (Cluster ii0 and ii3), 541 vascular tissue cells (Cluster ii2 and ii8), 207 epidermis 1 cells (Cluster ii6 andii7), 345 epidermis 2 cells (Cluster ii4), 659 subepidermis and shoot primordia cells (Cluster ii2), and 99 outgrowth shoot cells (Cluster ii9). In addition, Cluster ii5 was annotated as inner callus 2 based on its spatial location and lack of correlation with Cluster i (Figure 1D). The same was done for spots in Cluster iii and they were assigned to cell types including 145 inner callus 1 cells (Cluster iii6), 174 inner callus 2 cells (Cluster iii5), 139 inner callus 3 cells (Cluster iii8), 628 vascular tissue cells (Cluster iii2, iii3 and iii4), 139 subepdiermis cells (Cluster iii9), 277 epidermis 1 cells (Cluster iii0), 112 epidermis 2 cells(Cluster iii10), and 203 outgrowth shoot cells (Cluster iii7 and iii11) (Figure 1E). The classification was supported by the expression of marker genes (Supplemental Figure 1).

In addition, single-nucleus transcriptome profiling of tomato callus was conducted at the same developmental stage and a total of 17,214 single-nucleus transcriptomes with coverage of 23,909 genes using the 10x Chromium Single Cell. The 17,214 cells were grouped into 22 distinct clusters (Cluster iv, Supplemental Table 4) through principal component analysis and unsupervised analyses. These clusters were annotated based on their spatial classification features (Supplemental Figure 1), defined as inner callus1 (Cluster iv4 and iv10), inner callus 2 (Cluster iv12), vascular tissue (Cluster iv3, iv6, iv8, iv15 and iv19), Epidermis 1 (Cluster iv1, iv11 and iv13), Epidermis 2 (Cluster iv5) shoot primordia (Cluster iv9 and iv20) and outgrowth shoot (Cluster iv7) (Figure 1F).

### Cell differentiation from regenerated shoot primordia to outgrowth shoot

The samples were collected when a visibly regenerated shoot and a new shoot-meristem-like structure emerged from the tomato callus (Figure 2A). Further histological analysis showed that dense cells were the main morphological feature of the regenerated shoot primordia (Figure 2B and Supplemental Figure 2). Gene expression analysis, using markers homologous to those found in *Arabidopsis* shoot primordia (*PLT7, CUC1, WUS*, and *STM*), confirmed the cytological and molecular characteristics of shoot primordia in Sample i, ii, and iii (Figure 2C-2E). In addition, we identified new markers that are specifically expressed in shoot primordia cells of all samples (Supplemental Table 5), such as *COLD SHOCK DOMAIN PROTEIN 4* (*CSP4*), *HEAT SHOCK PROTEIN 70* (*HSP70)*, and *ARGONAUTE 4* (*AGO4*) (Supplemental Table 5), suggesting their potential roles in regulating tomato DNSO.

**Figure 2.**
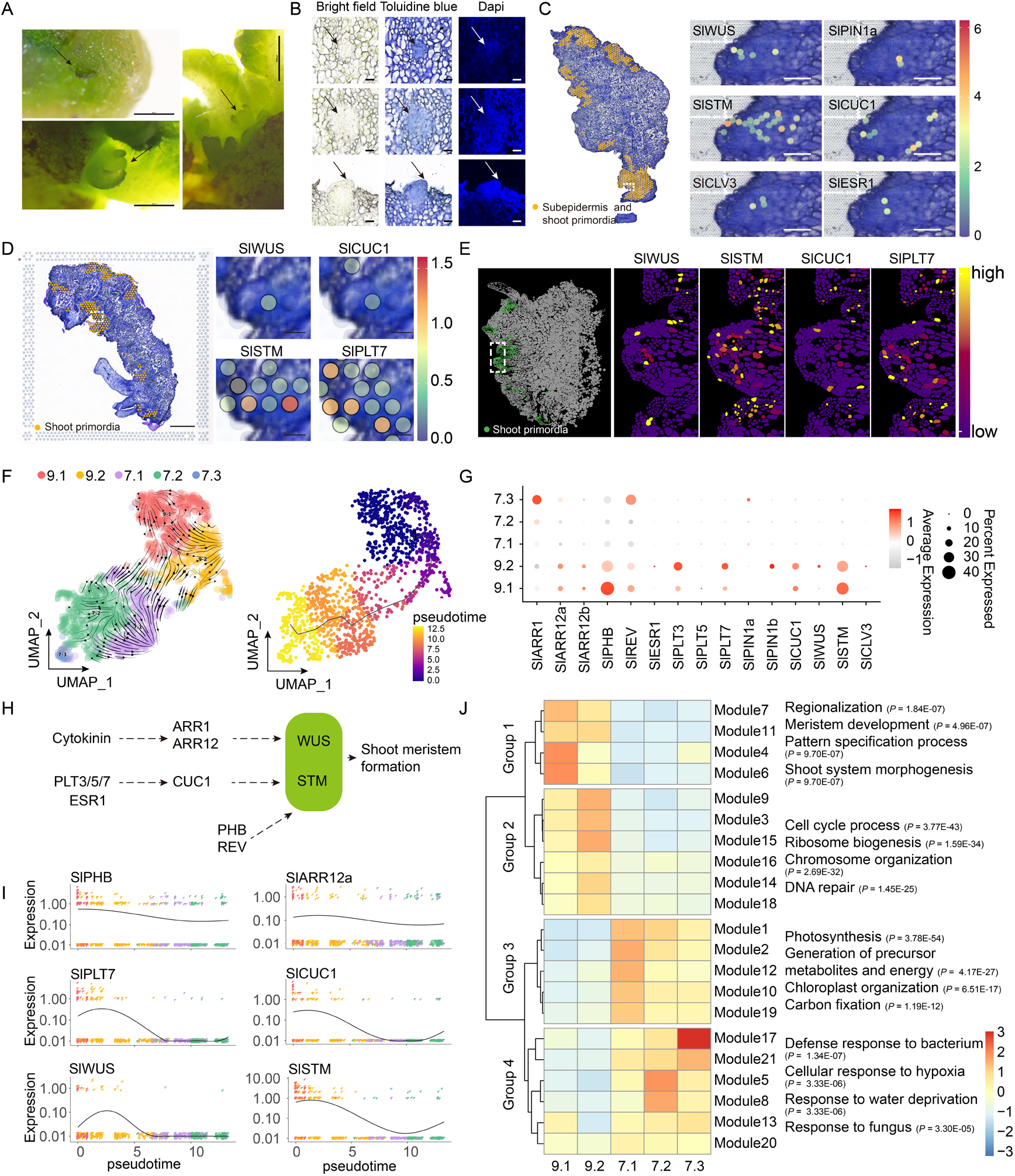
Cell Differentiation from Regenerated Shoot Primordia to Outgrowth Shoot. **(A)** Emerging regenerated shoot phenotypes with *de novo* shoot meristems indicated by arrows. The scale bar represents 1000 μm. **(B)** Histological analysis of shoot primordia showing the highly divided cell masses indicated by arrows. Tissue sections with 10 μm thickness were treated by TB and DAPI for cytological observations and nuclear staining, followed by image capture. The scale bar represents 50 μm. **(C-E)** Spatial gene expression of shoot primordium markers on Sample ii (C), Sample iii (D), and Sample i (E). Shoot primordium cells in samples were marked by the corresponding colors, and the spatial gene expressions of markers were enlarged and shown in the selected shoot primordia areas indicated by the dotted lines. **(F)** RNA velocity and pseudotime trajectory analysis of extracted shoot primordium cluster (9) and outgrowth shoot cluster (7) from Cluster iv. Sub-clusters (9-1, 9-2) and (7-1, 7-2, 7-3) are represented. **(G)** Dot plot of gene expression related to shoot regeneration in (F). **(H)** Predicted molecular regulatory pathway for shoot meristem formation in tomato callus. **(I)** Expression of typical shoot primordium promoters along the pseudotime axis. **(J)** Heat map of differentially expressed gene modules and functional enrichment in clusters 9-1, 9-2, 7-1, 7-2, and 7-3. The detailed GO enrichment information is listed in Supplemental Table 8.

To study the cell differentiation from shoot primordia to outgrowth shoot, cells defined as shoot primordia (Cluster iv9) and outgrowth shoot (Cluster iv7) from single-nuclear transcriptome were re-clustered. RNA velocity and Monocle3 pseudotime analysis revealed the cell fate decision from divided shoot primordia to differentiated shoots. The results supported a putative lineage transition along the 9.1-9.2-7.1-7.2-7.3 path (Figure 2F and Supplemental Figure 2). We conducted gene co-expression module analysis along the vector-field-based pseudotime axis and identified 21 modules grouped into four categories. The expression of genes in Group 1 decreased gradually from Cluster 9.1 to 7.3, and their enriched GO terms were related to regionalization, meristem development, and pattern specification, suggesting that cell fate was determined. Group 2 genes were enriched in Cluster 9.2 and related to the cell cycle, ribosome biogenesis, and chromosome organization, indicating active cell division. Group 3 and Group 4 genes were enriched in highly differentiated cells, and their GO terms were related to photosynthesis, precursor generation, defense response to bacteria, and cellular response to hypoxia (Figure 2J and Supplemental Table 8).

The expression of *SlWUS*, a key marker of shoot regeneration from callus, gradually increased from sub-cluster 9.1 to 9.2 within the shoot primordiaum (Figure 2C-2E and Supplemental Table 7). *SlSTM*, which helps maintain undifferentiated stem cells and directly interacts with *WUS* (Su et al., 2020), was more widely distributed in bud primordia than *SlWUS* and reached its peak expression level earlier on the pseudotime axis (Figure 2I). *SlCLV3*, activated by *WUS* to provide robustness for the *WUS* gradient (Plong et al., 2021), was restricted to a small spatial region during shoot regeneration (Figure 2C). *SlWUS*-expressing cells are usually located near the cytokinin response domain. B-type cytokinin response regulators (ARR1, ARR10 and ARR12) directly activate *WUS* expression to promote *Arabidopsis* shoot regeneration (Xie et al., 2018), with *SlARR12a* and *SlARR12b* being constant in tomato shoot primordia and *SlARR1* having a higher expression level in the late stage of the tomato outgrowth shoot. The expression of the *CUC* genes, which stimulates the growth of adventitious shoots when over-expressed (Takada et al., 2001), plays a crucial role in the formation of shoot primordia and is located upstream of *WUS* expression (Gordon et al., 2007). The level of *SlCUC1* expression remained high in Clusters 9.1 and 9.2 (Figure 2G, 2I), and its spatial location partially overlapped with that of *SlWUS* (Figure 2C-2E). The genes *ESR1, PLT3, PLT5*, and *PLT7*, which act as upstream regulators of *CUC1*, play a role in promoting shoot regeneration (Banno et al., 2001; Kareem et al., 2015). The expression levels of these genes, particularly *SlPLT7*, were high in the shoot primordia domain, with particularly high enrichment in Cluster 9.2. Additionally, Class III homeodomain-leucine zipper (HD Zip III) transcription factors, *PHB* and *REV*, are crucial for shoot regeneration and interact with *type-B ARRs* (Zhang et al., 2017) and activate *STM* expression (Shi et al., 2016). The expression levels of *SlPHB* and *SlREV* were higher in Cluster 9.1 than in Cluster 9.2, and their expression patterns on the pseudotime axis were similar to those of *SlARR12a*. In conclusion, *SlARR12, SlCUC1, SlPHB*, and *SlREV* are key regulatory factors in shoot regeneration from tomato callus and are involved in multiple pathways.

### Tomato Vascular Tissue in Callus Forms a Complex Nutrient Transport Network

Observations of frozen-sectioned callus under microscopy revealed vascular tissue formation in tomato callus, with distinct features. The vessel elements of the vascular tissue were observed in the culture medium after one day of explant placement (Figure 3Ai, left), and their enrichment was noted at the wound of the explant after three days of culture (Figure 3Ai right and Supplemental Figure 3). Callus contained abundant vascular tissue at the outgrowth shoot phase of shoot regeneration. Vascular tissue with xylem, procambial/cambium, phloem, and separated vessel components was observed, which were integrated into callus parenchyma cells and had diverse orientations (Figure 3Aii, 3Aiii and Supplemental Figure 3). Vascular tissue in different plant organs and growth stages has varying structures. In tomato callus, two main types were observed. One type had the xylem as its core, similar to that found in old roots or stems (Figure 3Aiii left and Supplemental Figure 3), while the other type had the xylem along the side, resembling leaf venation (Figure 3Aiii right). These types were arranged in multiple, branched strands in the callus, forming a complex vascular tissue network (Figure 3Aiv and Supplemental Figure 3). In the late phase, the vascular tissue of the outgrowth shoot connected to the network within the callus (Figure 3Av left and Supplemental Figure 3). A portion of the vascular tissue was adjacent to the culture medium (Figure 3Av right and Supplemental Figure 3), which may be related to the transport of nutrients from the culture medium to the front end.

**Figure 3.**
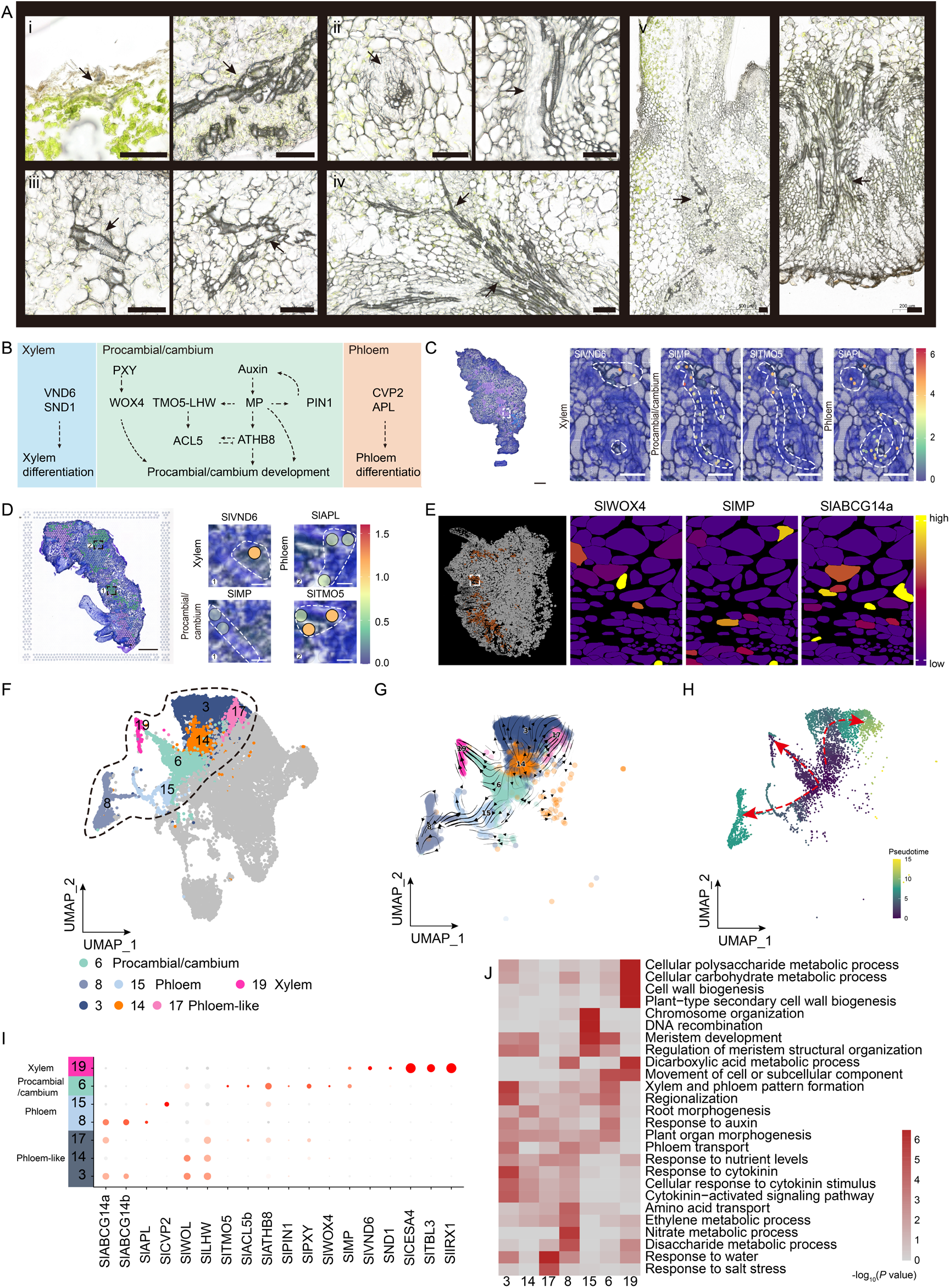
Vascular Tissue of Callus Forms a Complex Nutrient Transportation Network. **(A)** Cytological characteristics of vascular tissue as observed through histological analysis, with arrows indicating differentiated tissue. The scale bar represents 1000 μm. i, The vessel elements of the vascular tissue were observed in a culture medium after one day; ii, Enrichment of the vessel elements was noted at the wound of the explant after three days; iii and iv, The vessel elements were integrated with parenchyma cells and formed branched strands. v, The vascular tissue in the callus was connected to the shoot and located near the medium. **(B)** A predicted molecular regulatory pathway for vascular tissue development. **(C-E)** Spatial distribution and expression of key vascular tissue markers in Sample ii (C), Sample iii (D), and Sample i (E). The vascular tissue in the samples was marked by the corresponding colors, and spatial gene expression of markers was enlarged and shown in the selected vascular tissue areas indicated by the dotted lines. **(F)** Visualize the vascular tissue in Sample iv using UMAP, with different cell types marked by colors. **(G)** RNA velocity analysis of the vascular tissue clusters in (F), with red arrows indicating the predicted differentiation routes. **(H)** RNA pseudotime trajectory analysis of the vascular tissue clusters in (F). **(I)** Dot plot depicting the expression of genes in (B). **(J)** Enriched GO terms for the vascular tissue clusters in (F), with detailed information in Supplemental Table 8.

The role of vascular tissue in callus development and shoot regeneration is to provide delivery paths and mechanical support. Vascular patterning is regulated by complex genetic networks that vary in different plant organs and growth stages. The genetic basis for vascular tissue specification in callus has been established by examining of known genetic networks and their expression levels in spatial transcriptomes (Figure 3B). The main regulators of xylem vessel formation in the activated secondary cell wall are *VASCULAR-RELATED NAC-DOMAIN 6* (*VND6*) and *NAC SECONDARY WALL THICKENING PROMOTING FACTOR1* (*NST1*) (Kubo et al., 2005; Zhong et al., 2006). *SlVND6* expression was observed in sclerenchymatous cells through deep staining in frozen sections (Figure 3C and 3D). The auxin-induced *MONOPTEROS* (*MP*) is crucial for developing the procambium in the embryo and leaf vascular tissue (Hardtke and Berleth, 1998; Przemeck et al., 1996). The *MP* target gene, *TARGET OF MONOPTEROS 5* (*TMO5*), is sufficient to suppress the vascular initiation defects in *mp* mutants (De Rybel et al., 2013). *SlMP* and *SlTMO5* were expressed in parenchyma cells and arranged linearly (Figure 3C and 3D). *ALTERED PHLOEM DEVELOPMENT* (*APL*) is indispensable in promoting sieve element differentiation and suppressing xylem formation (Bonke et al., 2003). *SlAPL* expression was observed in regions close to procambial/cambium, suggesting active phloem development.

Vascular tissue cells that were derived from a single-nuclear transcriptome based on marker genes were further classified into various categories, including the xylem (represented by Cluster 19), the procambial/cambium (represented by Cluster 6), the phloem (represented by Clusters 8 and 15), and the phloem-like (represented by Clusters 3, 14, and 17) (Figure 3F). RNA velocity and Monocle3 pseudotime analysis indicated that the xylem and phloem originated from the procambial/cambium cells (Figure 3G and 3H). *SlMP, SlTOM5, ACAULIS5b* (*SlACL5b*), *ARABIDOPSIS THALIANA HOMEOBOX* 8 (*SlATHB8*), *SlPIN1a* and *SlWOX4* were enriched in the procambial/cambium cells, and the enriched GO terms for procambial/cambium genes were related to the formation of xylem and phloem, regionalization, response to auxin, and plant organ morphogenesis (Figure 3J). Representative genes involved in secondary cell wall development, including *VND6, SECONDARY WALL–ASSOCIATED NAC DOMAIN PROTEIN* (*SND1*), *cellulose synthase A4* (CESA4), *TRICHOME BIREFRINGENCE-LIKE 3* (*TBL3*) and *IRREGULAR XYLEM 1* (*IRX1*), were highly enriched in xylem cells. The presence of *COTYLEDON VASCULAR PATTERN 2* (*CVP2*), which regulates phloem specification via the control of phosphoinositide levels (Carland and Nelson, 2009), and *ARABIDOPSIS LOSER OF PHEROMONE RESPONSE 2* (*APL*), which plays a role in nuclear degradation during later stages of phloem development (Furuta et al., 2014), was observed in Cluster 15 and Cluster 8, respectively (Figure 3I). According to RNA velocity and pseudotime analysis, the cells in Cluster 15 were positioned at the initial stage of phloem development. The Cluster 15 showed an enrichment of GO terms related to chromosome organization and DNA recombination. On the other hand, Cluster 8 had an enrichment of GO terms related to amino acid transport, ethylene metabolism, nitrate metabolism, and disaccharide metabolism. Phloem-specific genes *ABCG14a, ABCG14b* and *WOODEN LEG* (*WOL*), related to cytokinin (Ko et al., 2014; Mähönen et al., 2006), were enriched in Clusters 3, 14 and 17, and defined as phloem-like cells. Interestingly, phloem-like cells were not rich in genes related to phloem development (Figure 3I), but genes in cytokinin–activated signaling pathway, response to nutrient levels, and plant organ morphogenesis were enriched. It was observed that the GO term related to cytokinin response was enriched in Cluster 3, while GO terms related to water and salt stress response was enriched in Cluster 17. These results suggested that phloem development in callus is active and diverse, and a rapid nutrient transportation network has been constructed. In summary, our findings suggest that the existence of specialized vascular tissue systems may re-conceptualize the callus as a temporary but comprehensive tissue, instead of the previously assumed uniform or multiple layers of cells, during plant regeneration.

### The Origin of Shoot Primordia and Its Spatial Alternative Splicing

The origin of shoot primordia or shoot progenitors is crucial in DNSO. While in *Arabidopsis* a subpopulation of pluripotent callus cells in the middle layer of the callus have been identified as shoot progenitors (Zhai and Xu, 2021), the same is not yet resolved in tomato. The single-cell resolution of Stereo-seq makes the cell types identified in Sample i more accurately reflect the biological state and clearly distinguishes them through cell boundaries. Interestingly, the shoot primordia cells were found to be tightly embedded in the sub-epidermis in the spatial clustering result of Sample i (Figure 4A), and the UMAP visualization also showed a close relationship between the two cell types (Figure 4B). These findings suggest that the shoot primordia may originate from the sub-epidermis, which was supported by further pseudotime trajectory analysis of both the shoot primordia and sub-epidermis (Figure 4C). However, more histological evidence is needed to verify this assumption.

**Figure 4.**
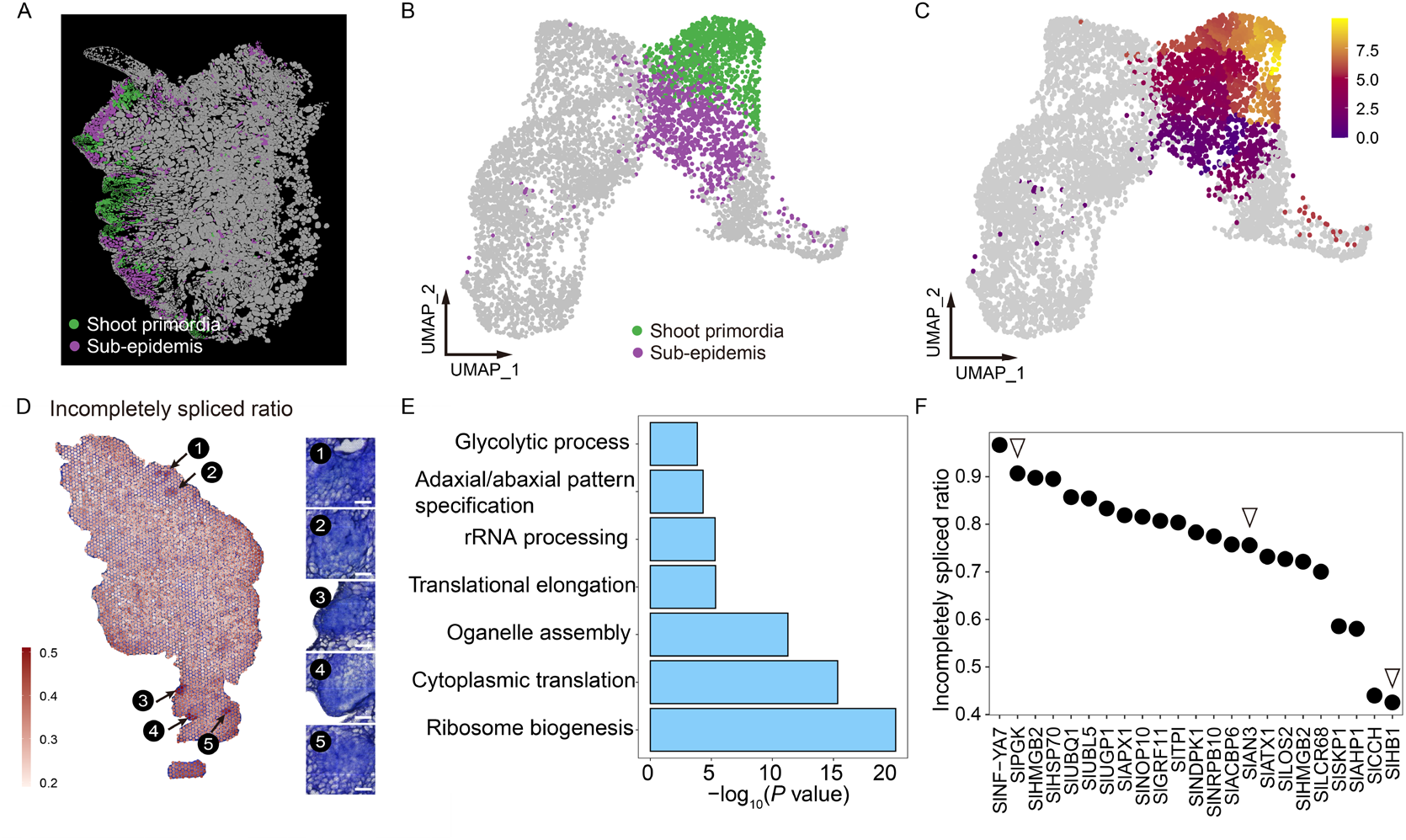
The Origin and Spatial Alternative Splicing of Shoot Primordia. **(A)** The single-cell level spatial distribution of shoot primordium and sub-epidermis in Sample i, marked by colors. **(B)** The UMAP visualization of shoot primordium and sub-epidermis in Sample i. **(C)** The pseudotime trajectory analysis of shoot primordium and sub-epidermis. **(D)** High IS ratio (> 0.4) and high expression (> 5 Normalized counts) of five regions in the shoot primordium are selected and shown with corresponding histological observations. The scale bar represents 100 μm. **(E)** The GO enrichment analysis for the genes selected from (D), with detailed information in Supplemental Tabel 8. **(F)** The IS ratio of the selected genes from (D), with *SlPG3, SlAN3*, and *SlHB1* were marked by inverted triangles.

We choose the BMKMANU S1000 platform for conducting ONT Nanopore long-read sequencing. As a result, we sequenced the Sample ii spatial transcriptome library again using the ONT PromethION 48. The average number of UMI and genes in 3,511 spots from the ONT data was 1,049 and 702, respectively. Comparison of UMI and gene counts between Illumina and ONT data revealed high consistency across spots (Supplemental Figure 4A). The long-read nanopore data provides information about transcript abundance and splicing isoform levels. As a result, we calculated the incompletely spliced (IS) ratio per spot, which had an average value of 31.6%. The results showed that the distribution of gene splicing was uneven, with the epidermis 1 (Cluster ii6 and ii7) and outgrowth shoot (Cluster ii9) regions exhibiting higher IS ratios (Supplemental Figure 4B). Furthermore, the shoot primordia (Cluster ii1) have a particular region with a high incompletely spliced (IS) ratio (Figure 4D). By analyzing 30 spots with an IS ratio of over 0.4 within the shoot primordia Cluster ii1, we discovered that genes with high expression (> 5 Normalized counts) and high IS ratio (> 0.4) were enriched in processes associated with ribosome biogenesis, cytoplasmic translation, translational elongation, and glycolysis (Figure 4E and Supplemental Table 6), pointing to a high level of protein synthesis activity in this area. Among genes with a high IS ratio (Figure 4F), we identified the glycolytic enzyme *PHOSPHOGLYCERATE KINASE* (*PGK*) (Rosa-Téllez et al., 2018), which suggests its potential role in providing energy for the shoot primordia region. *ANGUSTIFOLIA 3* (*SlAN3*) (Shimano et al., 2018) and *HEMOGLOBIN1* (*SlHB1*) (Miguel et al., 2020), which are related to leaf primordia initiation and leaf development, were also present, indicating that the fate of leaf primordia precursor cells was determined early during shoot regeneration.

## Discussion

The DNSO process in callus involves several stages of somatic cell reprogramming, cell fate determination, transformation, and development, indicating the presence of high callus heterogeneity. A prior single-cell RNA-seq study has found that *Arabidopsis* callus comprises three layers - inner, middle, and outer - and that pluripotency acquisition in the middle layer is essential for organ regeneration (Zhai and Xu, 2021). The activation of *WOX5*, in conjunction with its interaction factors *PLT1* and *PLT2*, synergistically promotes pluripotency acquisition in cells. In contrast to the focus on *Arabidopsis* callus initiate factor, our research utilized spatial and single-nucleus transcriptome profiling on tomato callus when significant shoot growth was observed. This revealed that the tomato callus composes multiple cell types, including epidermis, inner callus, vascular tissue, shoot primordia, and outgrowth shoot (Figure 1), thereby confirming the high cell heterogeneity of tomato callus at the molecular level.

The shoot primordia cells or shoot progenitors, presenting dense histological characteristics, are vital cell types during DNSO. Multiple genes in tomato with homology to key *Arabidopsis* shoot regeneration regulators, including *SlPIN1, SlARR1, SlARR12, SlPLT3/5/7, SlESR1, SlCUC1, SlPHB, SlREV, SlWUS*, and *SlSTM*, were detected in the shoot primordia region of callus through spatial and single-cell analysis (Figure 2), illustrating the conserved shoot regeneration regulation networks across species. The presence of one copy of the *SIWUS* gene in tomato, similar to *Arabidopsis*, implies its crucial role in tomato DNSO. However, the presence of multiple members of genes such as *SIPIN1a, SIPIN1b* and only one member of *SICUC* in tomato suggests the existence of distinct regulatory strategies in tomato DNSO. Additionally, it was found that *SlCSP4, SlHSP70*, and *SlAGO4* were identified in shoot primordium cells of all samples (Supplemental Table 5). The cold shock domain proteins play a role in developmental transitions in both plants and animals, and that overexpression of *CSP4* affects the late stage of embryonic development in plants (Yang and Karlson, 2011). The presence of *SlCSP4* in shoot primordia suggests its potential involvement in the transition of cell fate from shoot primordia to the outgrowth of a shoot. The high expression of *SlAGO4*, a crucial component in RNA-directed DNA methylation (Lahmy et al., 2016), in shoot primordium cells indicates that it could play a vital role in defining cell fate in that area. Furthermore, the presence of a high level of *SlHSP70*, a molecular chaperone known for its contribution to programmed cell death and plant immunity (Kim and Hwang, 2015), has also been noted. This protein is responsible for protein folding and remodeling processes and is often over-expressed in cancer cells in animals (Rosenzweig et al., 2019). Based on these observations, we infer that *SlHSP70* could be involved in regulating cell proliferation by altering the structure of target proteins during DNA synthesis. However, a deeper understanding of the molecular mechanism is necessary.

Previous histological studies have documented the formation of vascular tissue in callus during plant regeneration (Sangduen and Klamsomboon, 2001), but its development and potential functions have been overlooked. Our histological analysis revealed the diversity of vascular tissue in tomato callus and its partially interconnected distribution (Figure 3). Detailed cell type classification including xylem, procambial/cambium, phloem, and phloem-like cells, gene functional annotation and developmental trajectory were analyzed based on single-cell transcriptome. The results showed that amino acid transportation and nitrate metabolism were primarily carried out in phloem cells, suggesting that phloem cells play a role in nutrient transportation during tomato DNSO. Additionally, a type of phloem-like cell related to cytokinin was identified, where the expression of the cytokinin transporter, *ABCG14* (Zhang et al., 2014; Zhao et al., 2022), and cytokinin receptor, *WOL* (Kubiasová et al., 2020), was enriched. This indicates that the cytokinin-specific phloem-like cells are likely involved in efficiently transporting cytokinin from the external medium to the internal callus.

The application of spatial biology in plants presents both technical challenges and opportunities. One major challenge is the size difference among cells in the same tissue section, which can affect the efficiency of transcriptome capture. Fine slicing and longer permeabilization time are necessary for smaller cells, while the opposite is true for larger cells. To address this, the slice thickness and permeabilization time must be optimized based on cell size and research topic. Additionally, tissue cells with high water content, such as young roots, fresh stems, and emerging leaves, are prone to breaking during frozen sectioning, leading to RNA diffusion and transcript dislocation. Other challenges include transferring tissue sections to spatial slides, low UMI detection, AI-based cell boundary recognition, and precise scanning and matching of images. While the plant cell wall provides an advantage over animal research in demonstrating cell boundaries, it also negatively affects permeabilization and RNA release, which can be addressed by adding a cell wall digest step. Even though most spatial biology technologies are designed for animal sample studies, overcoming these technical challenges and advancing the field of plant spatial biology have great potential to lead to significant biological discoveries.

## Methods

### Plant Materials and Culture Conditions

Tomato (*Solanum lycopersicum* cultivar Micro-Tom) seeds were sterilized with 75% alcohol (2 min) and 3% sodium hypochlorite (15 min), then transferred to half-strength (½) Murashige and Skoog (MS) basal medium (2.22g/L MS, 10g/L sucrose, 3.5g/L phytagel, pH 5.8) and grown at 25° C conditions for 7 days. The cotyledon explants were sliced into 0.5 cm segments and cultured in medium (4.43 g/L MS basal medium with vitamins, 30 g/L sucrose, 1 mg/L ZT, 0.1 mg/L IAA, 3.5 g/L phytagel and pH 5.8) under 16-hour/8-hour light/dark conditions at 25 ° C for 16 days. Tomato regeneration reached the shoot outgrowth phase at 16 days, originated from callus. Samples were selected and applied to subsequent experiments.

### Tissue Cryo Sectioning

Fresh calli samples were placed into optimal cutting temperature compound (OCT, Sakura finetek Europe B.V.), and then vacuumed on ice for 5 min and centrifuged at 4 °C, 1000*g* for 5 min. The samples were subsequently transferred into the precooled embedding box filled with OCT. The embedding box and samples were fixed at -70 ° C, and then stored at -80 ° C. The pre-frozen samples in OCT were sectioned at 10 μm in thickness for spatial transcriptome sequencing and histological observation.

### Single-nucleus RNA-seq

Tomato callus with shoot outgrowth was selected for single-nucleus RNA-seq after removing residual explants. The fresh callus samples were cut with blade and dissolved in 1 ml extraction buffer (0.44 M SUCROSE, 1.25% Ficoll, 2.5% DXTRAN T40, 20 mM HEPES, 10 MM MgCL_2_, 0.5% TRITON X-100, 1 mM dithiothreitol (DTT), 1× protease inhibitor, 0.4 U/μl RNase inhibitor). The nuclear solution was filtered through 40 μm and 20 μm cell strainer into a new tube to remove tissue fragments, and then pelleted at 4 °C, 500*g* for 5 min. The pellet was washed once with 4 ml buffer, and then resuspended in 100 μl 1×PBS solution. The nuclei were stained with 4,6-diamino-2-phenylindole (DAPI) and counted with a cytometer. The final concentration was diluted to 1000-1200 nuclei/μl. The snRNA-seq library were constructed with 10x Genomic protocol (Chromium Single Cell 3’ Reagent Kits v3.1User Guide) and sequenced by Illumina NovaSeq.

The raw data were mapped to the tomato reference genome (version SL4.0 and Annotation ITAG4.1) by Cell Ranger 6.1.2 using default parameters. Filter matrix obtained was used for subsequent analysis by Seurat (v4.3.0) (Hao et al., 2021). Genes expressed in at least 3 cells and cells expressing at least 200 genes were initially selected. The matrix was further filtered using parameters such as nCount within twofold standard deviation, nFeature within twofold standard deviation, mitochondria percentage below 5% and chloroplast below 10%. The CellCycleScoring and SCTransform in Seurat were performed successively for matrix normalization and transformation. The first 30 dimensions obtained by RunPCA were applied to the analysis of RunUMAP, FindNeighbors, and FindClusters with 0.8 resolution to obtain UMP and cluster information.

### BGI Single-cell Stereo-seq

The scStereo-seq processes were performed according to the previously reported method with minor modifications(Xia et al., 2022). Briefly, tissue sections were adhered to the Stereo-seq chip surface and incubated at 37° C for 1 min, followed by the fixation in methanol for 30 min at -20° C. Then the tissue sections were treated with Fluorescent Brightener 28 (FB) for cell wall staining and with Qubit ssDNA HS Reagent for nuclear staining. Imaging was performed with a Motic CustomPA53 FS6 microscope. Permeabilization, reverse transcription, tissue removal, cDNA release and library preparation were successively performed according to the previous method. The spatial library was sequenced on a DNBSEQ-T10 sequencer.

The spatial representation matrix was generated from raw data according to the previously reported Stereo-seq method (Xia et al., 2022). The reads were then aligned to tomato reference genome using STAR (Dobin et al., 2013) and mapped reads with MAPQ 10 were counted and annotated to their corresponding genes using a script (available at https://github.com/BGIResearch/SAW). The cell was extracted based on the cell boundary information in the cell wall staining image, and the X-Y coordinates and transcript count of every single cell were obtained using customized R script. Finally, the spatial expression matrix at single-cell level was applied to the subsequent analysis. The R package, Seurat (v4.1.0), was employed for quality control, SCT normalization, dimensionality reduction, clustering, and identification of marker genes. To remove low-quality cells, cells with no more than 50 genes were filtered. After filtering, the remaining 6759 single cells with 21531 genes were included in the downstream analysis. Cluster analysis for the section was performed by FindClusters in Seurat (v4.1.0) with 0.5 resolution. Clustering results were displayed by uniform manifold approximation and projection (UMAP) dimension reduction analysis. Furthermore, spatial visualization of cell clusters and gene expression on cell mask were performed by customized R scripts.

### 10x Visium Spatial RNA-seq

At this stage, Visium Spatial Gene Expression Slide & Reagent Kit was employed. The 10 μm tissue sections were adhered to the chip surface and then fixed with methanol according to the method mentioned above. Then the tissue sections were stained with TopuidineBlue for cell wall staining and the imaging was captured with microscope slide scanner (PANNORAMIC MIDI II, 3DHISTECH Ltd). Permeabilization was performed firstly with treatment in RNase-free mix containing 10 mM DTT, 2% PVP, 0.2% BSA, 0.4% RNase inhibitor for 20 minutes at 37° C. Secondary permeabilization (9 min) and library construction were performed according to the user guide. The library was sequenced by Illumina NovaSeq.

The raw data were mapped to the tomato reference genome by Space Ranger 2.0.0 using the default parameters. The obtained filter matrix was used for subsequent analysis by Seurat (v4.3.0) (Hao et al., 2021). The first 15 dimensions obtained by RunPCA were applied to the analysis of RunUMAP, FindNeighbors and FindClusters with 0.6 resolution to obtain UMP and cluster information.

### BMKMANU S1000 RNA-seq

The MKMANU S1000 RNA-seq was performed with BMKMANU S1000 Gene Expression kit (BMKMANU, ST03002). Tissue sectioning, TopuidineBlue staining, imaging and first permeabilization were consistent with the user guide of BMKMANU S1000 Tissue Optimization Kit (BMKMANU, ST03003). Secondary permeabilization (9 min) and library construction were also performed according to the user guide. The Illumina library was sequenced with Illumina NavoSeq. In addition, some cDNA templates were adopted to construct the Nanopore library and sequenced with PromethION48.

The raw data from Illumina and Nanopore were mapped to the tomato reference genome by BSTMatrix v2.0 and BSTMatrix-ONT v1.1, respectively, using default parameters. The image adjusted by BSTViewer V1.42 and corresponding level 7 (50 μm) matrix were used for downstream analysis. The UMAP and clustering information were obtained using the R script (Bmk Space mapping.R, http://www.bmkmanu.com/archives/513) based on Seurat 4.3.0 (Hao et al., 2021) provided by the manufacturer (parameters: min.cells = 5, min.features = 100,. dims = 1:25, resolution = 0.5). The gene incomplete splicing ratio matrix was constructed using the sequences containing introns extracted by bedtools (Quinlan and Hall, 2010) and the file containing read id, barcode, gene id and UMI correspondence obtained by BSTMatrix..

### RNA Velocity Analysis

RNA velocity of single-nucleus transcriptome was evaluated by velocyto package (La Manno et al., 2018). The room file containing splicing information was generated with default parameters and read into ScVelo (Bergen et al., 2020). The UMAP and cluster information extracted by Seurat were also embedded in ScVelo. RNA velocity was estimated using full dynamic model, and projected to UMAP plot.

### Trajectory analysis

Pseudotime analysis for shoot regeneration and vascular development was performed using Monocle3 (Qiu et al., 2017). Graph-autocorrelation were analyzed through graph_test to find genes that vary over a trajectory, and then the modules of co-regulated genes were obtained through the find_gene_modules. Heat map of the average expression of genes with high variation in each module was generated according to the cluster. Genes with high variation were also subjected to GO biological analysis.

### GO enrichment analysis

Based on the known gene function and GO biological process in *Arabidopsis*, the homologs of *Arabidopsis* genes (TAIR10) in tomato were identified using BLASTP. GO enrichment analysis was performed using R package, clusterProfiler, with TAIR10 annotation as the background.

## AUTHOR CONTRIBUTIONS

B.L., and K.X. conceived and supervised the study. X.S., P.G., and L.C. performed bioinformatics analysis. M.W., J.Z., M.X., N.L., M.L., and L.F. conducted sampling and molecular lab experiments. B.L., K.X., X.S., and P.G. wrote the manuscript. B.L., K.X., X.X., and Y.G. participated in the manuscript editing and discussion.

## ACKNOWLEDGMENTS

We thank Wei Huang from BGI Shenzhen for sharing tomato seeds. We thank the 10x genomics headquarters at Pleasanton CA for the assistantce and critical comments on the subject. The authors declare that they have no competing interests.

